# hapCon: Estimating contamination of ancient genomes by copying from reference haplotypes

**DOI:** 10.1101/2021.12.20.473429

**Authors:** Yilei Huang, Harald Ringbauer

**Affiliations:** Department of Archaeogenetics, Max Planck Institute for Evolutionary Anthropology, Leipzig, Germany

**Author notes:** Corresponding author –.

## Abstract

**Motivation:** Human ancient DNA (aDNA) studies have surged in recent years, revolutionizing the study of the human past. Typically, aDNA is preserved poorly, making such data prone to contamination from other human DNA. Therefore, it is important to rule out substantial contamination before proceeding to downstream analysis. As most aDNA samples can only be sequenced to low coverages (<1x average depth), computational methods that can robustly estimate contamination in the low coverage regime are needed. However, the ultra low-coverage regime (0.1x and below) remains a challenging task for existing approaches.

**Results:** We present a new method to estimate contamination in aDNA for male individuals. It utilizes a Li&Stephen’s haplotype copying model for haploid X chromosomes, with mismatches modelled as genotyping error or contamination. We assessed an implementation of this new approach, hapCon, on simulated and down-sampled empirical aDNA data. Our results demonstrate that hapCon outperforms a commonly used tool for estimating male X contamination (ANGSD), with substantially lower variance and narrower confidence intervals, especially in the low coverage regime. We found that hapCon provides useful contamination estimates for coverages as low as 0.1x for SNP capture data (1240k) and 0.02x for whole genome sequencing data (WGS), substantially extending the coverage limit of previous male X chromosome based contamination estimation methods.

**Availability and Implementation:** A implementation of our software (hapCON) using Python and C has been deposited at https://github.com/hyl317/hapROH. We make hapCon available as part of a python package (hapROH), which is available at the Python Package Index (https://pypi.org/project/hapROH) and can be installed via pip. The documentation provides example use cases as blueprints for custom applications (https://haproh.readthedocs.io).

## Introduction

In recent years, ancient DNA (aDNA) has become a new powerful scientific instrument for studying the human past. However, aDNA is often highly fragmented and degraded, and the amount of endogenous DNA is typically low. Therefore aDNA is particularly prone to contamination from other human DNA, in particular during sample excavation, handling and aDNA extraction. It is an important quality control step to rule out substantial contamination before proceeding to downstream analysis. This task requires reliable tools to estimate contamination rates for low coverage aDNA.

One widely used approach to estimate contamination for aDNA utilizes heterozygosity in mitochondrial genomes (mtDNA) as an uncontaminated individual’s mtDNA should be haploid; therefore, apparent heterozygous sites on mtDNA contain evidence about contamination. [Renaud et al., 2015, e.g. Schmutzi]. For most ancient samples, mtDNA can be sequenced to relatively high coverage, facilitating such analysis. However, the ratio of preserved endogenous mtDNA to nuclear DNA varies greatly across samples, creating a complex relationship between mtDNA and nuclear DNA contamination. A sample can be highly contaminated for its nuclear DNA but minimally contaminated for its mtDNA, and vice versa [Furtwängler et al., 2018].

A direct way to estimate nuclear contamination similarly exploits the naturally haploid male X chromosome. Several methods have been developed to utilize this signal [Rasmussen et al., 2011, Moreno-Mayar et al., 2020, e.g.]. All of them require sites covered by at least two reads to measure heterozygosity. However, for low-coverage data most covered sites are covered by one read only. Assuming that read depth follows Poisson distribution per site, for 0.1x average genome-wide coverage about 0.47% sites are expected to be covered by at least 2 reads, for 0.05x dropping to 0.12% and for 0.01x to only 0.005%. As a result, only a small fraction of sequence data can be used for estimating contamination, causing the estimates for ultra-low coverage samples to be highly variable with wide confidence intervals.

Here we present a new approach to estimate contamination rates based on haplotype copying on male X chromosome that also utilizes sites covered by only one read. We model the target X chromosome as a mosaic copy from a modern reference panel, and model sporadic mismatches of observed reads from the copied haplotypes as either sequencing error or contamination. An implementation of the new method is available as Python package (hapCON, https://github.com/hyl317/hapROH). Using the Hidden Markov Model (HMM), the software estimates contamination by maximum likelihood. Extensive simulations and downsampling experiments demonstrate that hapCon produces estimates with smaller variance and narrower confidence intervals than previous methods using the male X chromosome. It substantially extends the application range of such analysis, yielding reliable contamination estimates for as low as 0.1x coverage on a widely used aDNA data type (1240k capture) and for as low as 0.02x whole genome sequencing (WGS) data (all coverages refer to average sequencing depth on the X chromosome).

## Methods

The core of our method is a haplotype copying approach widely used in genomics [Li and Stephens, 2003, Li&Stephen] that models a target genome as a mosaic of haplotypes from a reference panel. Since X chromosomes of males are haploid, they can be naturally modeled as such a haplotype mosaic[Biddanda et al., 2021]. Any reads discordant from the copied haplotype can be due to a number of causes (including mutation, gene conversion, sequencing error, aDNA postmortem damage, or contamination), but only contamination mismatches correlate with population allele frequencies. To utilize this signal within the HMM, we incorporated a previous contamination model for single sites [from ANGSD [Rasmussen et al., 2011]] which incorporates both mismatch rates due to contamination and due to other types of error.

### The Hidden Markov Model

Throughout, we model biallelic markers on haploid X chromosomes. Given *n* haplotypes from a reference haplotype panel, the Li&Stephens Hidden Markov model (HMM) has *n* hidden states for each marker, and a marker being in state *i* (1 ≤ *i* ≤*n*) denotes its genotype being copied from the reference haplotype indexed by *i* (Fig.1). This general Li&Stephen HMM is then fully specified by setting transition probabilities between markers and emission probabilities for the genotype data. Here we use a standard transition probability with jump probabilities depending on the genetic map distance between markers as measured in Morgan. For the emission probabilities of read counts for both alleles, we incorporate the previously published ANGSD model [Rasmussen et al., 2011].

**Figure 1:**
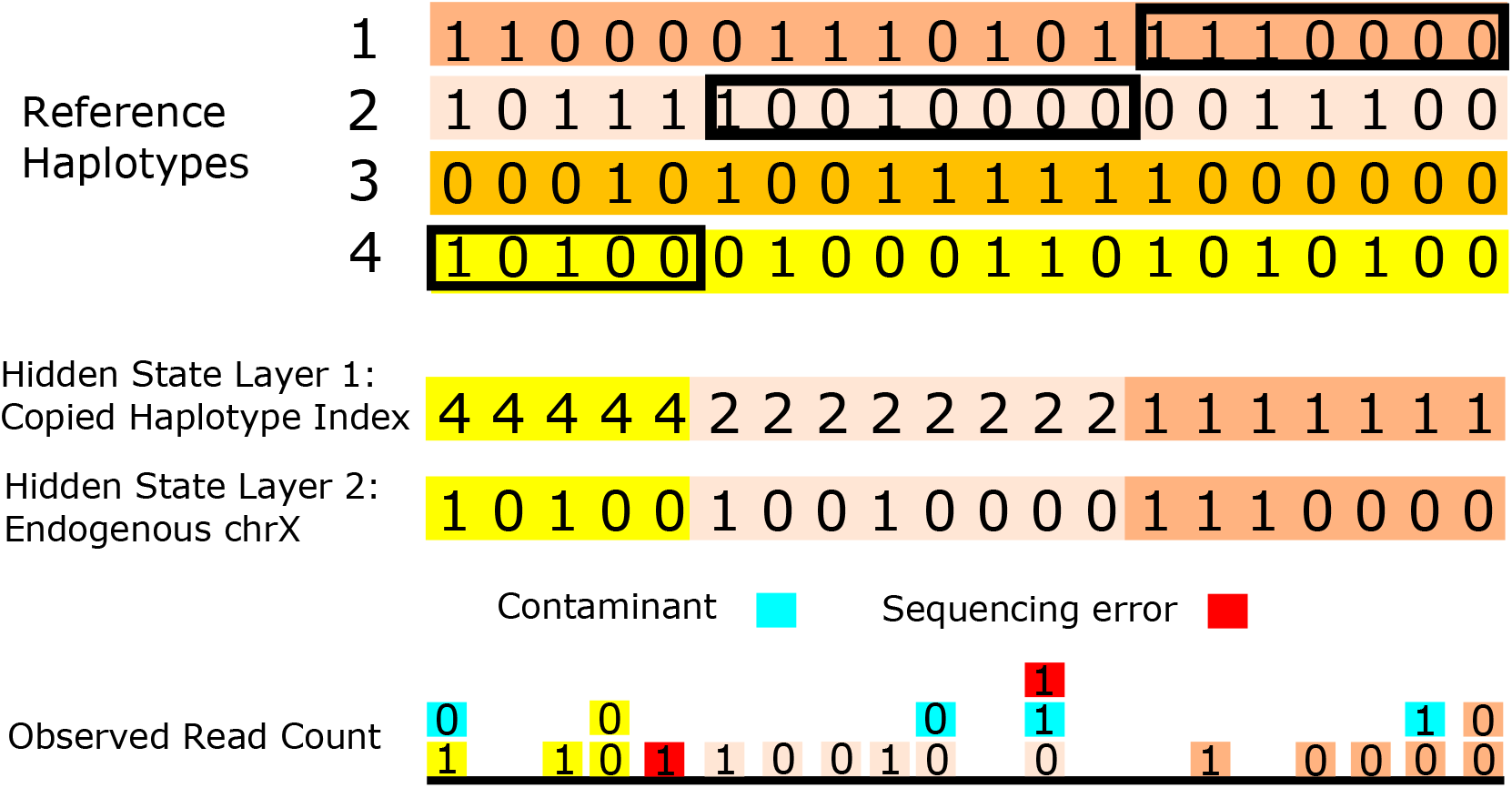
Graphical illustration of the model to estimate contamination rates via copying haplotypes from a haplotype reference panel. The target male X chromosome is modeled as a mosaic copy from a haplotype reference panel. In this specific case, the haplotype is copied from reference haplotype 4,2,1 (from left to right). The observed read counts at each biallelic marker are modelled as a mix of reads from the endogenous haplotype (shades of yellow), reads from the contaminant (blue) and sequencing errors (red).

### Transition Probabilities

We define the transition probability between hidden states for each pair of adjacent markers *l, l* +1 as in Ringbauer et al. [2021]. Given an infinitesimal rate matrix *Q* of dimension *n* x *n*, the full transition probability matrix between marker *l* and *l* + 1 is obtained by exponentiation of the rate matrix: *T*_*l*→*l*+1_ exp (*Q.r*_*l*_), where *r*_*l*_ denotes the genetic map distance between marker *l* and *l+* 1 (measured in Morgan). We assume that each reference haplotype has an equal prior probability to be copied from, therefore a single rate *q* fully specifies off-diagonal elements of *Q*, and *Q*_*ii*_ = -(*n* – 1) *q*. We set *q* “300, see Supplementary Note 1.3.3 for further details.

### Emission Probabilities

Assume we have known genotype data *i*_1_,*…, i*_*L*_ at *L* biallelic markers along the *i*th haplotype, with possible values 0 and 1 encoding reference and alternative alleles, respectively. At each marker *l*, we denote the number of aligned reads from the target sample supporting reference and alternative alleles by 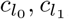, respectively. This observed read count data is the data we link to our HMM with our emission model.

To model mismatches between the observed genotype data and the copied haplotype, we now introduce three error parameters *ϵ*_*g*_, *ϵ*_*r*_, and *c*. First, *ϵ*_*g*_ is the genotyping error rate per read which can be estimated from monomorphic sites adjacent to polymorphic sites (see Supplementary Note 1.3.1 for details). Second, the overall error rate *ϵ*_*r*_ is an aggregate error term to model mismatches between the endogenous genotype and the copied haplotype due to various causes (including mutations, gene conversion, errors in the reference panel, etc.). This so called miscopying rate is widely used in phasing and imputation algorithms based on the Li&Stephen’s model [Loh et al., 2016, Delaneau et al., 2019, Rubinacci et al., 2021, Browning et al., 2021, e.g.]. We fix *ϵ*_*r*_ = 1*e*^-3^, as preliminary tests indicated that this value provides good performance on simulated and empirical aDNA data while also providing some flexibility so that the copying path is not truncated by errors (see Supplementary Note 1.3.2 for details). Third, the contamination rate *c* models the fraction of the reads originating from contamination, which is the parameter we wish to estimate.

We then use a two-layer approach to model the observed read counts of the target at each marker. The first layer models the endogenous genotype given the copying state, and the second layer describes how sequencing reads are drawn given the endogenous genotype.

The first layer specifies the genotype probability of the target haplotype for each marker *l, t*_*l*_, given the underlying copying state, *s*_*l*_. Haplotype copying with error rate *ϵ*_*r*_ gives:

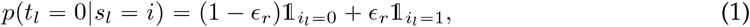

and the probability for the derived allele *p*(*t*_*l*_ *=* 1| *s*_*l*_ *= i*) is obtained analogously.

The second layer then models the probability of a read being derived given the latent genotype. We first calculate the probability of a single sampled read being derived. Let *c* denote the genome-wide contamination rate, and *p*_*l*_ denote the derived allele frequency in the contaminating population at marker *l*, then the probability of observing a derived read is

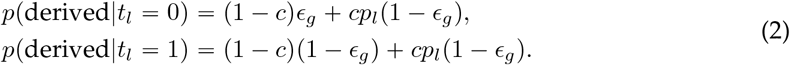

We have omitted terms that are proportional to *ϵ*_*g*_*c*, which are the products of two small values.

The probability for a single read being derived given the hidden state *s*_*l*_ is obtained by combining the two layers. Summing over the two possible latent genotypes gives:

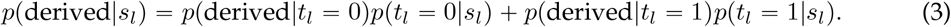

Finally, we model the observed read counts by a binomial distribution fully determined by the probability of a single read being derived. Denoting the total read depth at marker *l* as 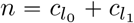 and abbreviating *p*_*d*_(*s*_*l*_) “*p*(derived|*s*_*l*_) gives:

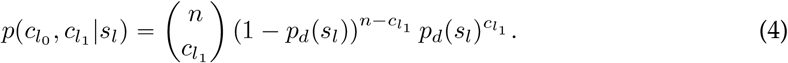

This probability of the observed read counts for each HMM state fully specifies the emission probabilities of the HMM.

### Maximum Likelihood Estimation

For a given contamination rate *c* and with all other parameters set, we then use a standard scaled forward algorithm to calculate the overall likelihood of the HMM model [Bishop, 2006]. To obtain a maximum likelihood estimate ĉ of *c*, we then use the iterative method L-BFGS-B[Byrd et al., 1995, Zhu et al., 1997] provided in SciPy [Virtanen et al., 2020] searching within the interval [0, 0.5]. We estimated the standard error of the MLE estimate ĉ by numerically calculating Fisher Information of the likelihood function around ĉ using the Python package numdifftools, and then approximated the 95% confidence interval by ±1.96 standard errors. Since our model is undefined for *c* <0, for ĉ = 0 the first derivative may not be zero at ĉ = 0 and thus confidence intervals cannot be approximated with the Fisher Information matrix alone. Instead, we use quadratic interpolation based on first and second derivatives with *c* to approximate the likelihood function around ĉ = 0 and use the set of parameters whose likelihood is at least 14.7% of the maximum likelihood to obtain 95% confidence intervals (the so-called “14.7% likelihood region”, see details in Rossi [2018], Definition 5.11).

### Relations to previous methods

Several methods that utilize discordant reads in haploid regions to estimate contamination rates in aDNA data have been developed. ANGSD, a widely used method, assumes that true endogenous allele is supported by the majority of the mapped reads at a site [Rasmussen et al., 2011]. More recently, a similar approach has been developed which assigns equal priors to both the reference and alternative alleles [Moreno-Mayar et al., 2020, two-consensus method]. Our new approach can be considered as a many-consensus model where the true endogenous allele originates from a set of reference haplotypes and each of them being weighted by the Li&Stephens haplotype copying framework that utilizes linkage disequilibrium information. We note that in the limit of widely spaced markers with no haplotype information left, our model converges to the two-consensus approach, but with priors according to the allele frequency in the reference panel.

We hypothesized that our method’s performance gain is driven by its ability to utilize sites covered by only one read. Such data can be used by neither ANGSD nor the two-consensus approach as both need at least two reads per site to establish evidence of contamination. In contrast, by using haplotype copying model, our method can detect potential contamination via comparing single reads to the copied reference haplotypes. As a proof of concept, we simulated read counts and down-sampled every covered site to exactly one read (see Supplementary Note 1.2 for details). Our results demonstrate that our method can still produce valid contamination estimates, even when fully relying on this so-called pseudohaploid data (Fig. S1).

## Results

We assessed the performance of our new approach on both simulated and empirical aDNA data. Throughout, we set the following default settings. We used a reference panel consisting of all non-African haplotypes from the 1000Genome Project[Consortium et al., 2015] (see section “Genetic distance between the endogenous and contaminant ancestry” for the detailed rationale). We set allele frequencies of the CEU individuals (CEU: Northern Europeans from Utah in 1000 Genome panel) as the proxy for the contamination source allele frequency. For comparison, we used ANGSD’s Method 1(new llh) with default settings. We filtered mapped reads to mapping quality greater than 30 and to base quality greater than 20. For each simulated scenario, we generated 100 independent replicates. For every replicate, we report the maximum likelihood point estimate of the contamination rate and a 95% confidence interval.

We prepared two reference panels tailored towards two common aDNA data types. The first panel contains all sites in the widely used enrichment capture strategy consisting of ca. 1.2 million SNPs, henceforth referred to as “1240k” panel [Fu et al., 2015, Haak et al., 2015, Mathieson et al., 2015]. The second panel contains all biallelic sites in the WGS 1000Genome dataset [Consortium et al., 2015] with minor allele frequency greater than 5%, henceforth referred to as “1000G panel”. We chose this 5% MAF filter because initial exploratory analysis showed that this cutoff provides a robust trade-off between accuracy and run time (Fig. S5). We explored MAF ranging from 0.2% to 20% and found that the width of confidence interval increases only slightly when increasing MAF cutoff, suggesting that most signal comes from common variants.

### Performance

#### Simulated whole genome sequencing data

We first assessed our new method on simulated samples with artificial contamination created by mixing BAM files of two samples. We chose WGS data with high average coverage and little contamination from the Allen Ancient Genome Diversity Project (https://reich.hms.harvard.edu/ancient-genome-diversity-project). Specifically, we used I1496 (5211-4958 calBCE, Hungary, contamination estimated by ANGSD: 0.756%(95% CI: 0.702%-0.810%))[Gamba et al., 2014] as the ancient source and I5319 (1050-1400 calCE, Alaska, USA, contamination estimated by ANGSD: 0.720%(95% CI: 0.665%-0.774%))[Flegontov et al., 2019] as the contaminant source. We randomly subsampled the original high-coverage BAM files and mixed them together to create synthetic BAM files with desired coverage and contamination rate. We then estimated contamination rates with both ANGSD and hapCon on those synthetic files.

We explored our method’s ability to distinguish between minimally and highly contaminated samples, a typical application in aDNA studies and present estimation results for 0% and 10% simulated contamination; results for other contamination rates (from 0% to 20% are visualized in Fig. S6 and Fig. S7). Comparing with ANGSD shows that our method produces estimates with smaller variance and narrower confidence intervals, particularly at lower coverages (Fig. 2). Overall, hapCON achieves a similar level of uncertainty as ANGSD at ca. 10x lower coverage when using the 1000G reference panel and at ca. 2x lower coverage when using the 1240k reference panel. In addition, we observe a marked boost in power when using the 1000G panel compared to the 1240k panel. With this panel, our method can robustly distinguish 10% contaminated samples from no contamination for as low as 0.02x X chromosome coverage (Fig. 3). We also observe a modest upward bias of our method at coverages higher than 0.5x when using 1000G panel. Further exploration shows that this bias remains mild at high coverages. (Fig. S9).

**Figure 2:**
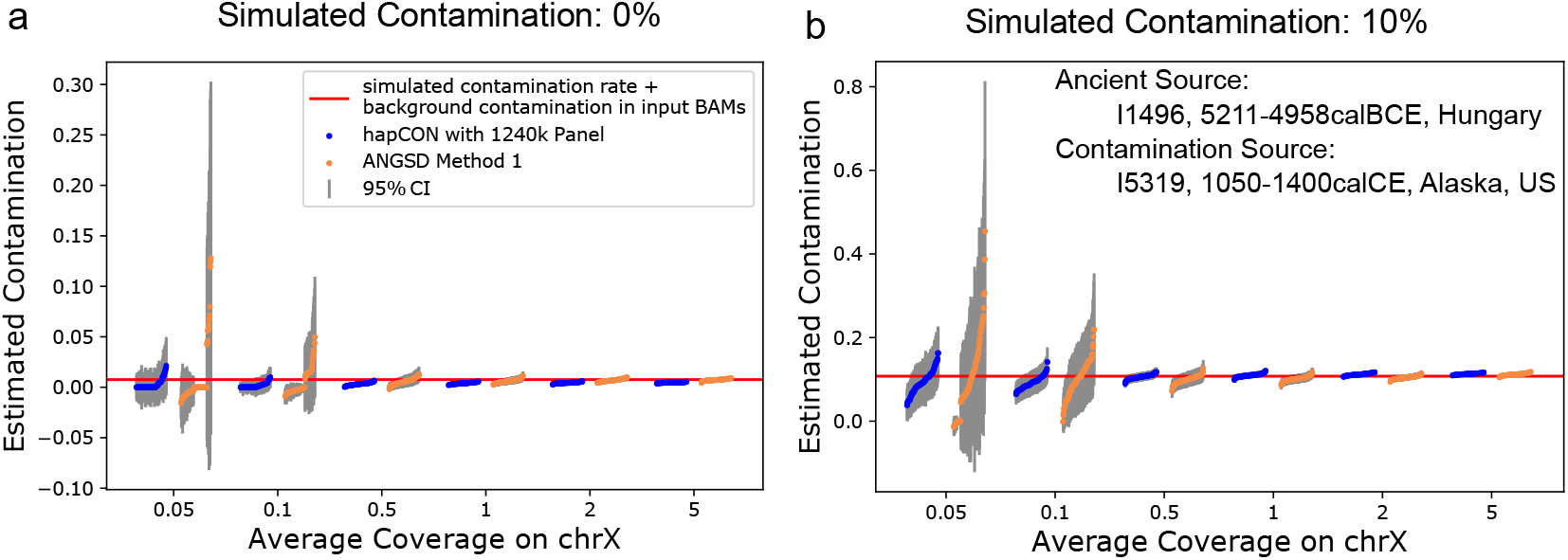
Performance on 1240k Panel for Simulated Contaminated BAM Files. We simulated contaminated BAM files by mixing two minimally contaminated BAM files (I1496, contamination: 0.756%(95% CI: 0.702%-0.810%); I5319, contamination: 0.720%(95% CI: 0.665%-0.774%)). 100 replicates were created for each simulation scenario and analyzed with hapCon and ANGSD. Estimates from both methods are visualized in groups of replicates next to each other. Each represents the estimate for one replicate, and they are ordered from low to high within each replicate group. Baseline contamination from I1496 was added to the simulated contamination rate (red line). Results for contamination rate other than 0% and 10% are displayed in Fig. S6. **a** Comparing ANGSD with our method using 1240k panel on simulated non-contaminated BAM files. **b** Same as (a) but with 10% simulated contamination.

**Figure 3:**
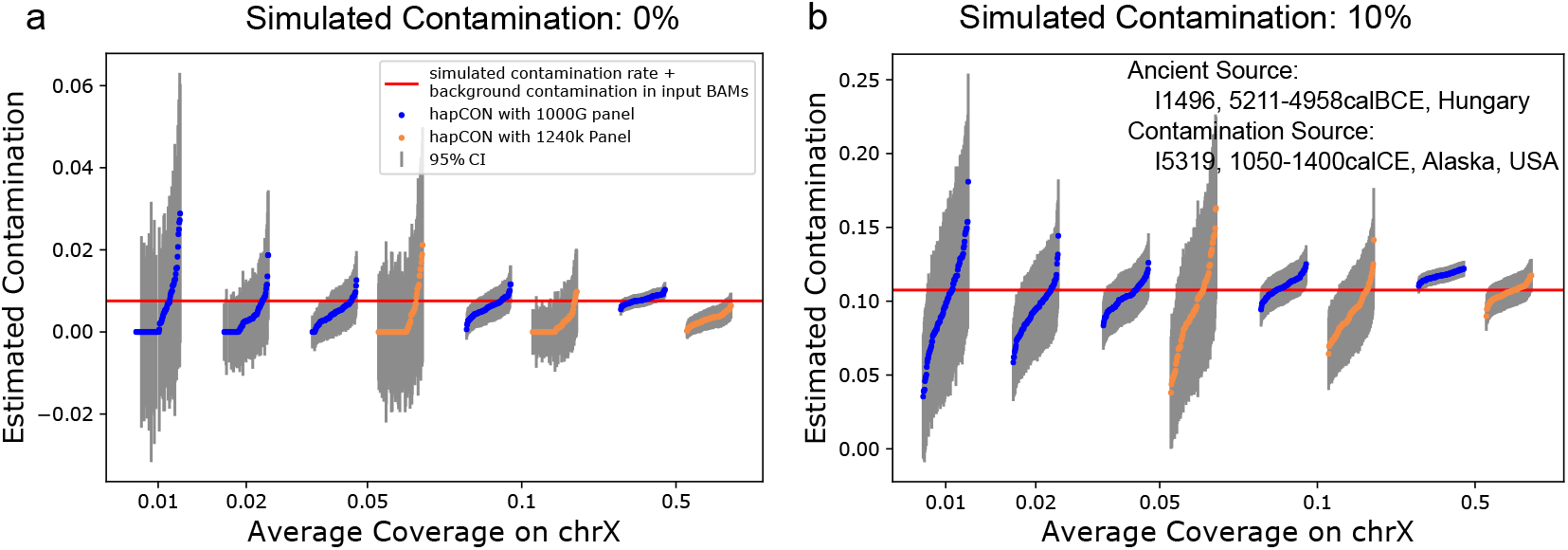
Performance on 1000G Panel for Simulated Contaminated BAM Files. We compare our method’s performance when using the 1240k and 1000G reference panel. Simulation settings are as described in Fig. 2. For coverages lower than 0.05x the estimates from the 1240k panel have very high variance, therefore we omitted them from the visualization. Results for contamination rates other than 0% and 10% are shown in Fig. S7. Baseline contamination from I1496 was added to the simulated contamination rate. **a** Comparing our method on two different reference panels for BAM files with no added contamination. **b** With 10% added contamination.

#### Down-sampling empirical aDNA data

We then tested the new method on published BAM files from previous aDNA studies when down-sampling genomic coverage. For 1240k data, we explored two male individuals from Sardinia, SUA001 (1411-1228 calBCE, 1.02x chrX coverage on 1240k target sites) and SUA002 (2274-2032 calBCE, 0.64x chrX coverage on those sites) [Marcus et al., 2020]. We chose those two samples because ANGSD estimates SUA001 to be substantially contaminated (10.45%, 95%CI: 9.56%-11.34%) and SUA002 to be only slightly contaminated (0.38%, 95%CI: 0.072%-0.69%).

For each target coverage, we independently down-sampled 100 replicates (Fig. 4). For the highly contaminated sample (SUA001) at coverage 0.05x, our method identifies 98 replicates as having substantial contamination (here defined as >5%), while ANGSD falsely identifies 20 replicates as being lowly contaminated (<5%). For the minimally contaminated sample (SUA002) at coverage ∼0.05x, our method can confidently identify all 100 replicates as minimally contaminated (<5%), while ANGSD’s estimate ranges from 0% to greater than 5%, falsely identifying two replicates as having substantial contamination. For 0.1x coverage, our new method can robustly distinguish minimally and substantially contaminated samples - all the down-sampled SUA001 replicates have contamination estimates greater than 5%, and all SUA002 replicates have contamination estimates less than 5%. Based on this down-sampling experiments, we recommend our method for 1240k data with 0.1x coverage or higher.

**Figure 4:**
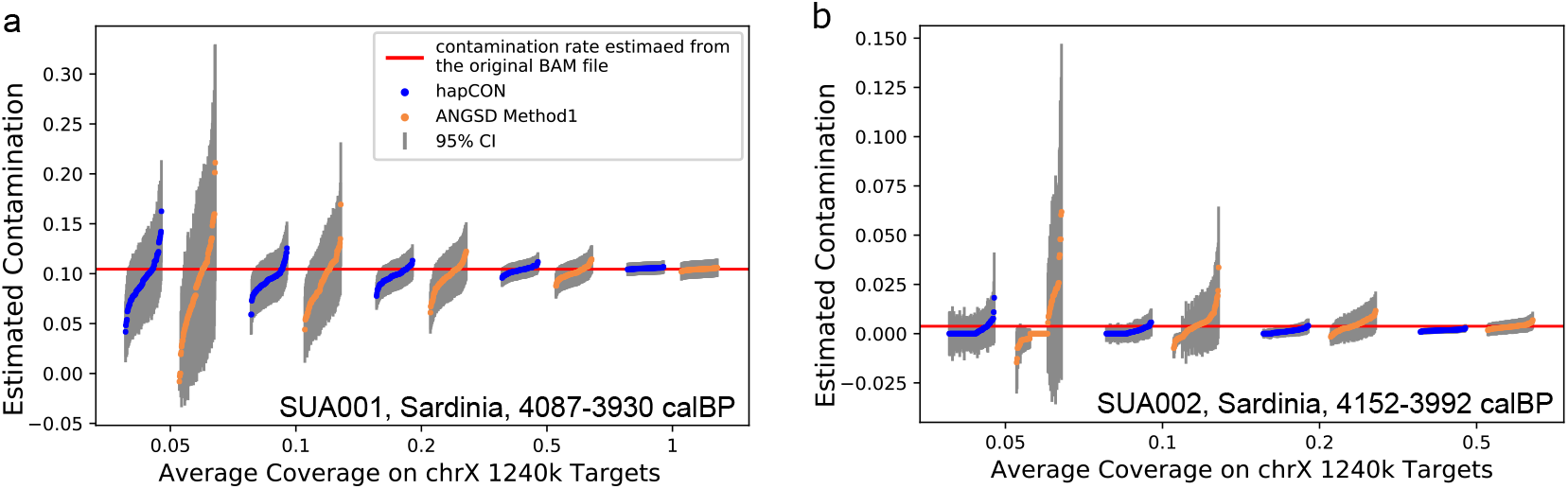
Down-sampling Experiment on 1240k Data of Two Sardinian Individuals. The original BAM files [Marcus et al., 2020] were down-sampled to various coverages, with 100 independent replicates for each target coverage. **a** Comparison between our method and ANGSD on SUA001, estimated to be 10.45%(95%CI: 9.56%-11.34%) contaminated by ANGSD (using the full data). **b** Comparison between our method and ANGSD on SUA002, estimated to be 0.38%(95%CI: 0.072%-0.69%) contaminated by ANGSD (on full data).

Further, we applied the two-consensus method [Moreno-Mayar et al., 2020] to these two Sardinian samples. We found that it performs overall similarly to ANGSD, but on some replicates much worse at 0.05x coverage (Fig. S8). Therefore, we focus our overall analysis on comparison between our new method and ANGSD, a currently very widely used method.

For down-sampling experiments of WGS data, we used a XiongNu sample DA43(Mongolia, 400BCE-100CE, 0.83x chrX coverage)[de Barros Damgaard et al., 2018], which is estimated to be 2.83% (95% CI: 2.35%-3.31%) contaminated by ANGSD. Since this is WGS data, we used the 1000G reference panel. We tested our method’s performance on coverage 0.01x, 0.02x, 0.05x, 0.1x, 0.5x and compared our method to ANGSD (Fig. 5). Our results show that our method yields reliable estimates for WGS data down to about 0.02x on X chromosome, outperforming ANGSD and achieving similar confidence intervals at 10x lower coverage, a performance gain similar to that observed on the simulated contamination BAM files.

**Figure 5:**
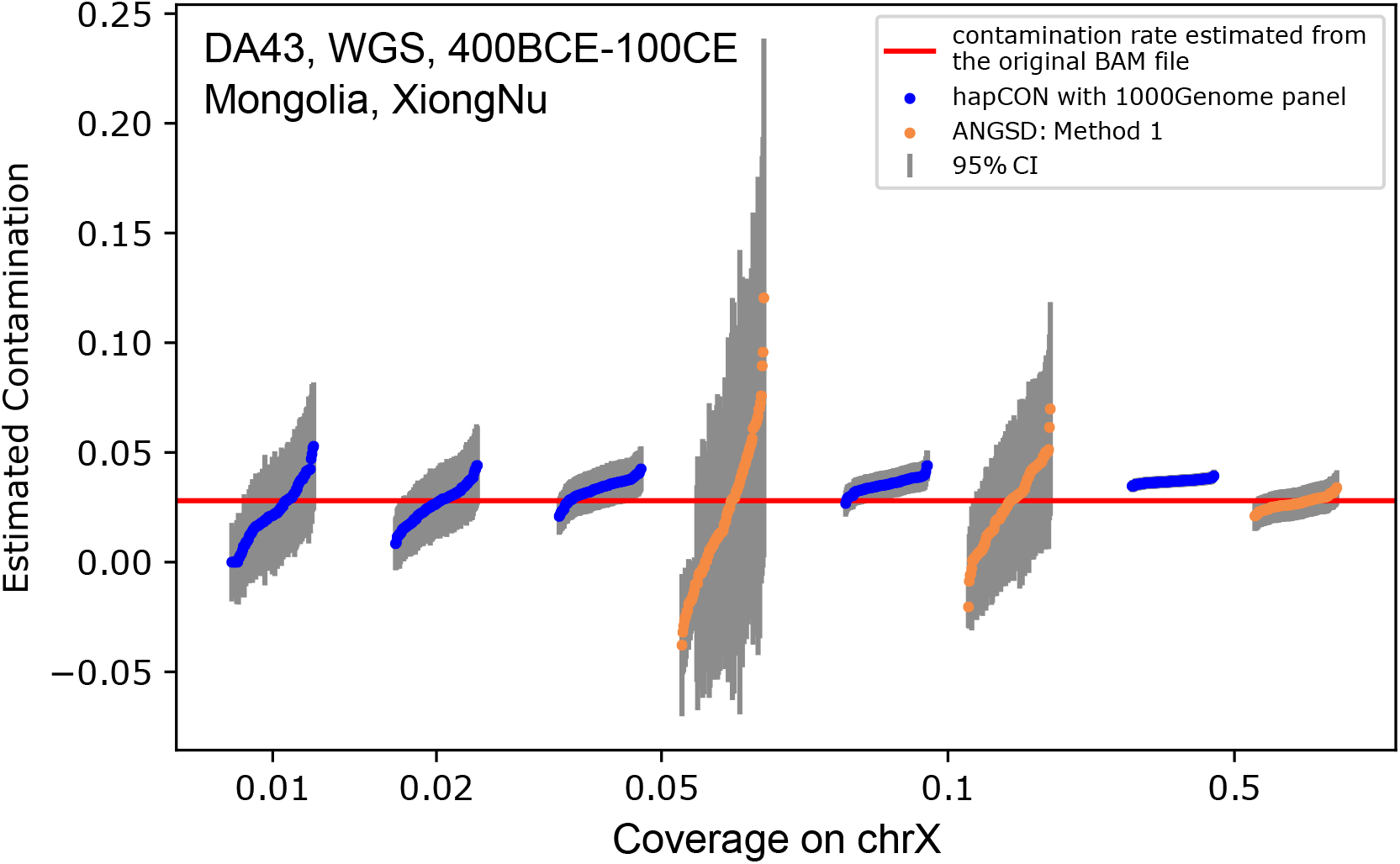
Down-sampling Experiment on WGS Data of DA43, XiongNu, Mongolia. The original BAM file for DA43 was down-sampled to various coverages 0.01-0.5x, with 100 independent replicates for each target coverage. We only visualized ANGSD’s results on 0.05x, 0.1x, 0.5x as it produced highly variable estimates at coverage lower than 0.05x. We depict the ANGSD contamination estimate (2.83%, 95% CI: 2.35%-3.31%) when using the full data (red line).

#### Empirical comparison on 1240k data

Next, we applied our method to empirical 1240k aDNA data covering a wide range of coverages and contamination rates. We selected all 89 ancient males that have X chromosome coverage greater than 0.05x from [Olalde et al., 2019], all of which are from the Iberian Peninsula and date to within the past 8,000 years. We also applied our method to 60 male Eurasian hunter-gatherer samples to test our method on older samples which are genetically more distant from the modern reference panel. We found that estimates from hapCON and ANGSD are highly concordant on the full sample set (*r*^2^ = 0.855), and for 139 out of 149 samples our method has smaller confidence intervals (Fig. 6). For the 119 samples with contamination rate estimated to be <5% by both methods, our the estimate of hapCon is higher than that of ANGSD on 38, and lower on the remaining 81, indicating that both methods give overall similar estimates when the contamination rate is low. For all samples with contamination rate greater than 20%, our method estimates higher contamination than ANGSD; however, in practice samples with contamination rate substantially greater than 10% are avoided in downstream analysis in any case.

**Figure 6:**
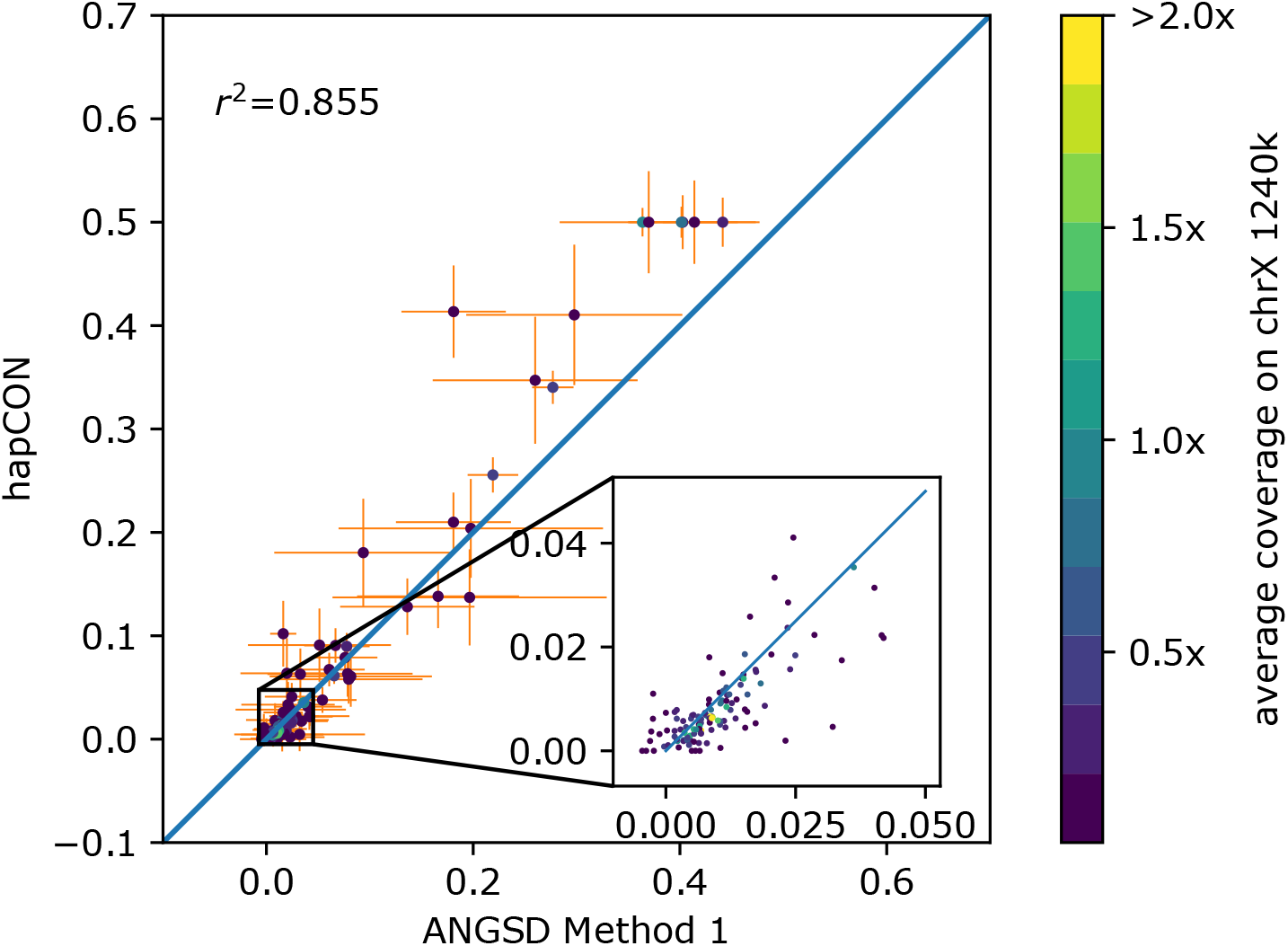
Comparison between the new method and ANGSD on 1240k aDNA data of 89 Iberians and of 60 Eurasian hunter-gatherers. The true contamination rate is unknown. No down-sampling is performed and all individuals (dots) are color coded by the average coverage on chrX 1240k target sites. The inlet shows a zoom-in into [0, 0.05] x [0, 0.05].

#### Implementation and runtime

We implemented hapCon as a Python package, expanding upon code from the software hapROH which uses a similar copying HMM [Ringbauer et al., 2021]. We measured our method’s runtime (including preprocessing time to parse BAM file with samtools [Li et al., 2009, Li, 2011]) on Intel(R) Xeon(R) Gold 6240 CPU @ 2.60GHz on WGS data with coverages ranging from 0.02x to 5x. As expected, the run time grows approximately linearly with the number of sites covered by at least one read. The run time of our method with 1000G panel remains within three minutes for a typical aDNA sample with coverage less than 1x, making our new method viable for any large-scale ancient DNA studies. Our benchmarking experiment also shows that our method is four times slower with 1000G panel than with 1240k panel (Fig. 7), as expected since the 1000G panel contains about four times more SNPs than the 1240k panel. For comparison, we used the C++ version of ANGSD. The results indicate that our method is faster than ANGSD at coverage higher than 1x.

**Figure 7:**
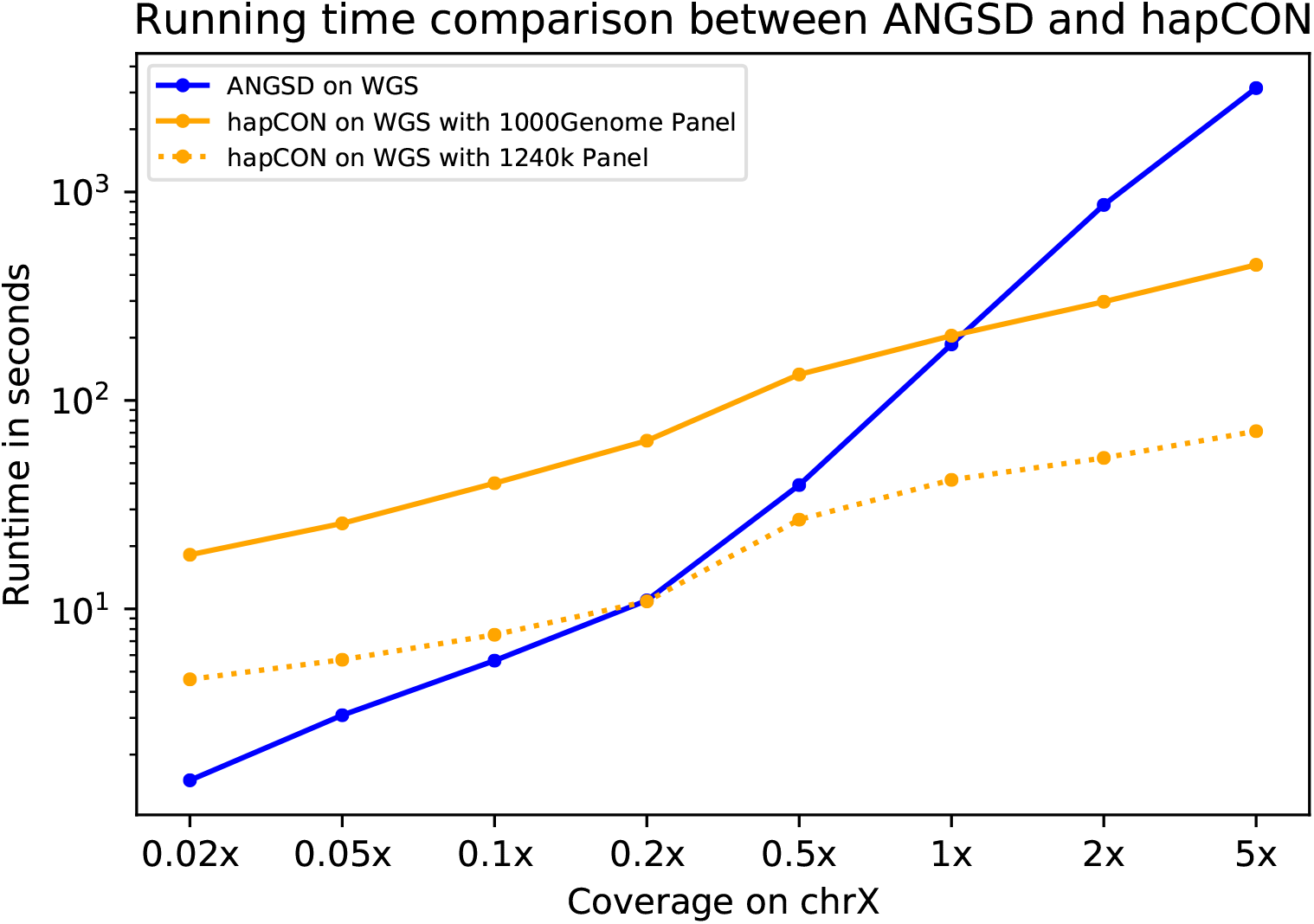
Comparing runtime of hapCON and ANGSD. We measured runtime of hapCON and ANGSD on BAM files of individual I1496, down-sampled to 8 target coverages. For hapCON, we used two different reference panels (1240k and 1000G panel). Each point represents the runtime averaged over 10 independent runs.

### Model Mis-specification

#### Mis-specified contaminant allele frequency

In practice it is often not possible to specify the ancestry of the contamination. One may not have an accurate proxy for the contamination source, or a sample may be contaminated by more than one sources of contamination. Therefore, a contamination estimation method is ideally robust to mis-specifying allele frequency of the contamination source.

To assess the effect of mis-specified contaminant allele frequency, we utilized synthetic BAM files simulated as described above. We then estimated contamination rate using allele frequencies from CEU (Utah residents with Northern and Western European ancestry), FIN (Finnish in Finland), GBR (British from England and Scotland), IBS (Iberian Populations in Spain), TSI (Toscani in Italia), YRI (Yoruba in Ibadan, Nigeria), CHB (Han Chinese in Beijing, China), PEL (Peruvian in Lima, Peru). We observe that the estimates obtained from CEU, FIN, GBR, IBS, TSI allele frequency behave very similar and all have little bias, while estimates using YRI allele frequency show significant downward bias (Fig. S11 and Fig. S12). We note that a similar downward bias for mis-specified contaminant ancestry was previously described for the two-consensus method [Moreno-Mayar et al., 2020]. These observations indicate that contamination estimates with our method remain robust with respect to modest allele frequency misspecification; however, mis-specification at the level of intercontinental allele frequency differences can introduce substantial biases. Our method provides a command line argument to specify the source of contamination; therefore, users can specify a contaminant ancestry that is different from the default.

#### Genetic distance between the endogenous and contaminant ancestry

We observed that, when the ancestry of contaminant sequences (e.g, CEU) is genetically close to that of endogenous sequences (e.g, TSI), our model tends to overestimate contamination rate at low coverage (0.05x, see Fig. S10a,d). We hypothesized that this bias is caused by the contaminant allele frequency being a better fit for the endogenous sequence than the reference panel overall. At such low coverage, almost every site is covered by only one read and covered sites are often far apart. Without haplotype structure, the main information for estimating contamination then comes from allele frequencies. When the allele frequency of the specified contamination source is closer to the endogenous haplotype than the allele frequency of the overall reference panel, there is a bias toward the contamination source. Indeed, when haplotypes of African ancestry are removed, the upward bias substantially decreases (Fig. S10b,e). When using the allele frequency calculated from the full reference panel, so that there is no allele frequency difference between the reference panel and the specified contaminant ancestry, the upward bias is completely removed (Fig. S10c,f). However, we observed that using global allele frequency creates downward bias at low coverages in empirical aDNA data (data not shown), plausibly because of overall allele frequency mis-specification of the contaminant. Therefore, we recommend using allele frequency as closely matching the true contamination source as possible and removing highly divergent haplotypes from the reference panel.

## Discussion

We have presented a new approach to estimate aDNA contamination in males based on a Li&Stephen’s haplotype copying model and implemented it in a software package (hapCon). The Li&Stephen model is widely used in population genomics as it makes use of haplotype structure and linkage disequilibrium information, and constitutes a central part of many modern phasing and imputation algorithms [Loh et al., 2016, Delaneau et al., 2019, Rubinacci et al., 2021, Browning et al., 2021, e.g.]. Similarly, our method implicitly imputes the endogenous genotype using reference haplotypes, and thus can utilize sites covered by only a single read, which, to our knowledge, cannot be utilized by any other male X chromosome based method. Testing on simulated and down-sampled empirical aDNA data showed that the new approach substantially improves power to estimate contamination, particularly in the low coverage regime. Across coverage levels, hapCON consistently yields estimates with lower variance and narrower confidence intervals than ANGSD and the two-consensus approach described in [Moreno-Mayar et al., 2020]. The most substantial gains are achieved for low-coverage WGS data. We found that hapCON provides robust contamination estimates for 1240k capture data with as low as 0.1x coverage and for WGS data with as low as 0.02x coverage on the male X chromosome, substantially extending the limits of ANGSD or the two-consensus approach. We explored various sources of model mis-specifications, including genotyping error, haplotype copying jump rate and contaminant allele frequencies. These experiments showed that hapCON is robust with respect to reasonable mis-specifications.

There are several methods to estimate contamination that do not rely on haploid regions, but they have limited applicability. ContamLD utilizes breakdown of linkage disequilibrium introduced by contaminant sequences to estimate contamination rate since the contaminant haplotype is uncorrelated with the endogenous haplotype[Nakatsuka et al., 2020]. As a key advantage it allows estimating autosomal contamination for female samples, but it requires comparably high coverage (∼0.5x for 1240k data and ∼0.1x for WGS data - which is about 5x higher than hapCon). Another recently introduced method, AuthentiCT, uses postmortem damage pattern to estimate contamination rate [Peyrégne and Peter, 2020]. It can work with very low coverage samples, but it is limited to single-stranded libraries without UDG treatment, limiting its usage to only a small fraction of aDNA data. Moreover, AuthentiCT cannot detect contamination originating from ancient sources. DICE performs joint estimate of demography, sequencing error and contamination rate but it requires comparably high coverage (∼ 3x), which is not obtained for the vast majority of ancient DNA samples[Racimo et al., 2016].

There are several limitations of our new approach. Haplotype copying substantially improves the power; however, it requires that the true endogenous haplotype can be modeled well as a mosaic of modern haplotypes. Deeply diverged human lineages such as Neanderthals and Denisovans are likely outside the range of this copying model. In such cases, one should consider using ANGSD or other more specialized methods not relying on a haplotype reference panel [Peter, 2020]. Having that said, we have tested our method on Paleolithic and Mesolithic hunter-gatherers and found good correlations between estimates from our method and that from ANGSD; therefore, this haplotype copying approach in principle work for most modern human aDNA. We also found moderate upward bias at low coverage when the endogenous and contaminant allele frequencies are genetically close. Our results indicate that this bias is caused by attraction to allele frequencies of the contamination source. Using a Out-of-Africa haplotype reference panel partially alleviates this bias. Finally, although our results showed that our method is in general robust to moderately mis-specified ancestry of contamination, to obtain unbiased results the allele frequency should be within continental genetic variation of the true contamination source. Therefore, if there is no prior information about the contamination source or the sample has been contaminated by several sources from different continental ancestries, our method may yield substantially biased results.

Beyond application to the naturally haploid male X chromosome, we envision our haplotype copying approach to be useful for estimating contamination for female samples with long runs of homozygosity (ROH), as such regions are effectively haploid. Previous studies have identified extensive ROH in almost all paleolithic hunter-gatherers[Ringbauer et al., 2021] or in populations with small effective size, such as the pre-contact Caribbean [Fernandes et al., 2021]. However, we note that contamination interferes with identifying ROH, particularly in the low coverage regime. Future work could establish robust approaches to identify ROH for substantially contaminated data, and the software presented here can then be straightforwardly extended for estimating contamination on ROH.

## Acknowledgements

We thank Yu He, Cosimo Posth and Johannes Krause for providing BAM files of hunter-gatherer samples prior to their publication.

## Funding and Competing Interests

This work was supported by funding from the Max Planck Society. The authors declare no competing interests.

## Author Contributions

We annotate author contributions using the CRediT Taxonomy labels (https://casrai.org/credit/). Where multiple individuals serve in the same role, the degree of contribution is specified as ‘lead’, ‘equal’, or ‘support’.

- Conceptualization (Design of study) – lead: HR; support: YH
- Software – YH; support: HR
- Formal Analysis – YH
- Data Curation – YH
- Visualization – YH
- Writing (original draft preparation) – lead: YH; support: HR
- Writing (review and editing) – YH, HR
- Supervision – HR
- Funding Acquisition – HR

## Data Availability

No new DNA data was generated for this study. The two Sardinian samples SUA001 and SUA002 are publicly available through the European Nucleotide Archive (ENA) under accession PRJEB35094. The Iberian samples are publicly available through ENA under accession PRJEB30874. The Mongolia XiongNu sample DA43 is available through ENA under accession PRJEB20658. The two high-coverage ancient genomes I1496 and I5319 are publicly available at Allen Ancient Genome Diversity Project https://reich.hms.harvard.edu. The collection of 60 hunter-gatherers is unpublished data and will become publicly available with the publication of this data (Yu et al., in preparation). The raw reference panel data that we used (phased haplotypes from the 1000 Genomes dataset, Phase 3, release 2013050) is available at http://ftp.1000genomes.ebi.ac.uk/vol1/ftp/release/20130502/. The reference panel needed to run our method is available at https://www.dropbox.com/sh/mxsf2c7srlx2qhm/AADVE5-gk5hp10nZjJw9fjMPa?dl=0.

## Code Availability

A implementation of our software (hapCON) using Python and C has been deposited at https://github.com/hyl317/hapROH. We make hapCon available as part of a python package (hapROH), which is available at the Python Package Index (https://pypi.org/project/hapROH/) and can be installed via pip. The documentation provides example use cases as blueprints for custom applications (https://haproh.readthedocs.io/en/latest/). A list of software and Python packages used in this work can be seen at Supplementary Section 2.

## Notes

https://reich.hms.harvard.edu/ancient-genome-diversity-project

### Competing Interest Statement

The authors have declared no competing interest.

