## Supplementary Materials for "hapCon: Estimating contamination of ancient genomes by copying from reference haplotypes"

December 2021

### 1 Additional Simulations

Several parameters in our model need to be set, but the ideal values for those can often not be exactly determined for each use case. To assess the general robustness of contamination estimates, we performed under the model simulations, where we first generate data as assumed in our generative HMM model and then evaluated our method using mis-specified parameters.

#### 1.1 Default Simulation Settings

Here we describe the default setting for the simulations. Single parameters are then altered around their default values to test our model under a variety of model mis-specifications, as described below.

To create genotype data, we first copy haplotype blocks with genotype error rate $1e^{-3}$ from TSI (Tuscany, Italy) haplotypes in the 1000Genome Dataset (Phase 3, [Consortium et al., 2015]). A copied haplotype block is chosen randomly with equal probability from all reference haplotypes, with each copied segment having length drawn from an exponential distribution with mean 1/3 centimorgan. At the end of each copied block, a new reference haplotype is chosen and copied from. Having simulated a mosaic of reference haplotypes, for each marker $i$ in the 1240k panel we then draw read counts from a Poisson distribution with mean equal to the target coverage multiplied by a weighing factor $\lambda_i$. This weighing factor models that in 1240k capture data some sites are systematically more likely to be covered than others. We obtain this weighting factor $\lambda_i$ by comparing site coverage to genome-wide average coverages in all male samples in Olalde et al. [2019]. Contaminant reads are drawn according to the global allele frequency in the 1000Genome dataset. To simulate read genotype error, we flipped the genotype of every read to the other allele with probability $1e^{-2}$. As reference panel for inference, we used all 1000Genome haplotypes excluding TSI samples.

#### 1.2 Downsampling Simulated Read Counts to Pseudohaploid Data

In one simulation scenario, we randomly sampled one read for each marker covered by at least one read. This procedure simulates how the so-called pseudohaploid data is generated from aDNA data. We note that, even though each site is only covered by one read, a high coverage sample will have more sites covered and therefore still contain more information than a low coverage sample. Our results show that our method produces robust contamination estimates even for such pseudohaploid data (Fig. S1). In practice, we recommend using our method on read counts directly generated from a BAM file to make full use of all information, but this simulation of pseudohaploid data demonstrates the power of utilizing haplotype structure for estimating contamination.

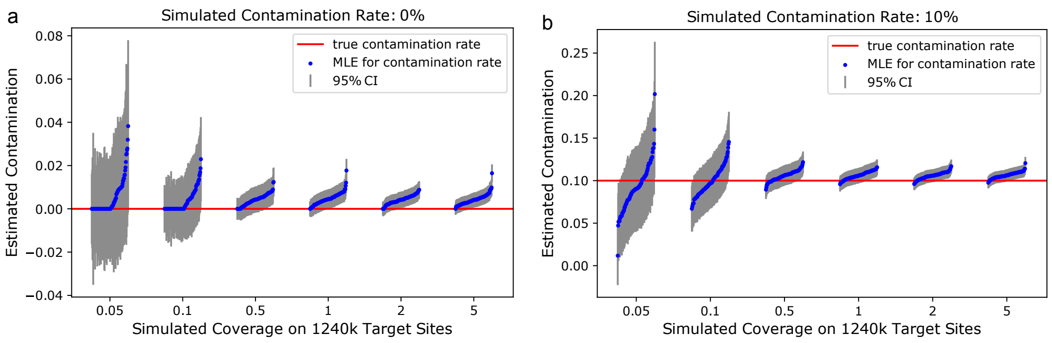


Figure S1: **Performance on Simulated Pseudohaploid Data at Various Coverages**

#### 1.3 Model Mis-specification

##### 1.3.1 Mis-specified Genotyping Error

In practice, we estimate genotyping error $\epsilon_g$ from discordant reads at sites flanking to the sites contained in the reference panel, as is done in Rasmussen et al. [2011], Moreno-Mayar et al. [2020]. These sites are expected to be invariable in the population; therefore, any discordant reads reflect only sequencing error and not contamination. Suppose there are a total of $L$ non-polymorphic sites in the reference panel and let $M_l, m_l$ denote the count of major and minor reads at site $l$ respectively. Then we estimate genotyping error $\epsilon_g$ by

$$\epsilon_g = \frac{\sum_{l=1}^{L} m_l}{\sum_{l=1}^{L} M_l + m_l}$$

This implicitly assumes that sites adjacent to markers in the reference panel are fixed, including the contamination source. We use four adjacent sites on either side of the polymorphic marker to estimate genotyping error.

When taking read counts from a BAM file, we only count reads matching either the reference or alternative allele in the reference panel. Reads that match neither of these two alleles are discarded. Therefore, we set $\frac{\epsilon_g}{3}$ as the genotyping error parameter in our HMM model since $\epsilon_g$ as estimated above captures the possibility of misreading a base to all three other bases. We note that this is an average approximation; for example, post-mortem damage preferentially produce C→T and A→G mismatches. However, explicit modeling of biased error rates is complex and case-dependent, here we aim for a robust approximation for a wide range of application scenarios.

To evaluate how mis-specified genotyping error rate affects our method, we simulated 10% contamination as described in Section 1.1 except that we varied the simulated sequencing error ranging from $1e^{-4}$ to $1e^{-2}$, evenly spaced on a log scale. We then estimated contamination rate when setting $\epsilon_g = 1e^{-3}$ for all simulated samples. This value is within the typical range of error rate estimates from empirical aDNA data.

We observe moderate upward bias of contamination estimates when the specified genotyping error rate is substantially below the true error rate (Fig. S2). A plausible intuitive reason for this bias is that if the mismatch observed cannot be fully explained by genotyping error, then the method attributes mismatches to contamination. Importantly, we observe that bias remains on the same order of magnitude as the mis-specification of the error rate. In empirical aDNA studies the error rate is usually on the order of $1e^{-3}$, whereas one wishes to estimate contamination on the order of $1e^{-2}$ or higher. Therefore, mis-specified genotyping error should not introduce relevant biases in most empirical analyses.

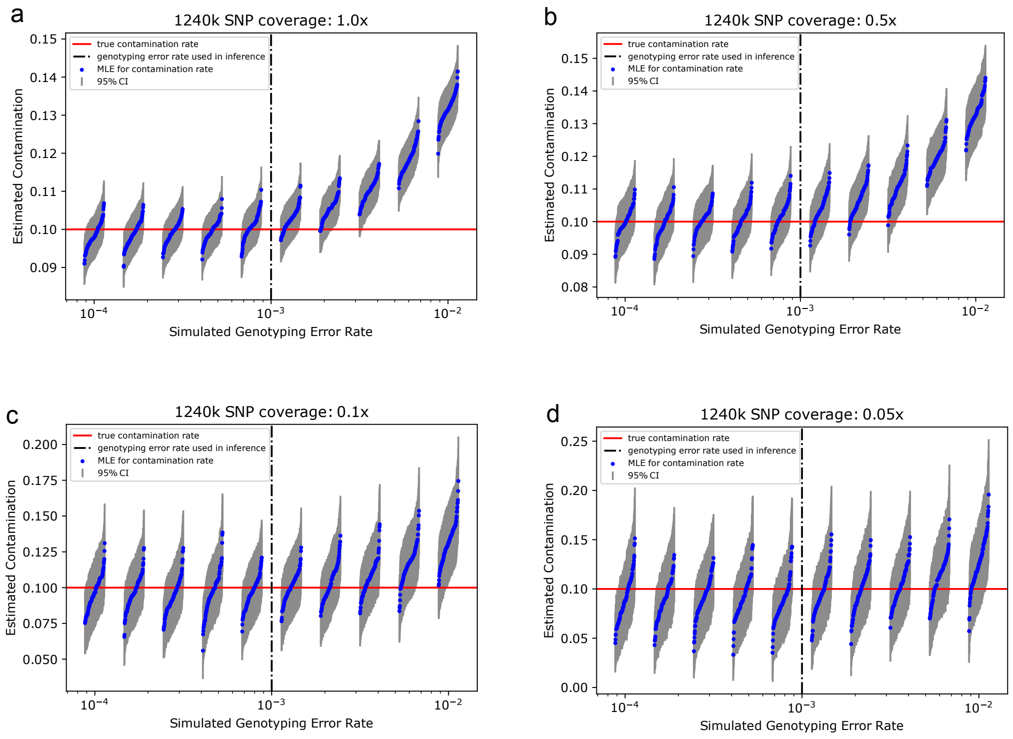


Figure S2: **Effect of Mis-specified Genotyping Error at Various Coverages**

##### 1.3.2 Mis-specified Haplotype Copying Error Rate

Some events such as mutations, gene conversions, or errors in the reference panel can cause sporadic genotype mismatches between the copied haplotype and the target haplotype. Therefore we use a copying error rate to model such mismatches so that a copying path would not be cut short prematurely. Following other methods that use Li&Stephen's copying model[Rubinacci et al., 2021, Loh et al., 2016], we set $1e^{-3}$ as the default copying error rate. To evaluate the effect of mis-specified copying error rate, we simulated 10% contamination as described in Section 1.1

except that we varied the miscopying rate from $1e^{-4}$ to $1e^{-2}$, evenly spaced on a log scale. We then estimated contamination rates when setting $\epsilon_r = 1e^{-3}$ for all simulated samples.

The results indicate that the contamination estimate is robust to mis-specified copying error rate (Fig. S3). Even in cases where the miscopying rate is one magnitude higher than assumed ($1e - 3$), upward bias remains minimal at 1.0x, 0.5x and 0.1x, and only becomes moderate at 0.05x. Similar to the case of mis-specified genotyping error discussed above, we observe that bias remains on the same order of magnitude as the mis-specification of mis-copying rate. In empirical data the error rate is usually on the order of $1e^{-3} - 1e^{-4}$, depending on the genetic distance between the endogenous haplotype and the modern haplotypes, whereas one wishes to estimate contamination on the order of $1e^{-2}$ or higher. Therefore, mis-specified mis-copying rate should not introduce substantial biases in empirical analyses.

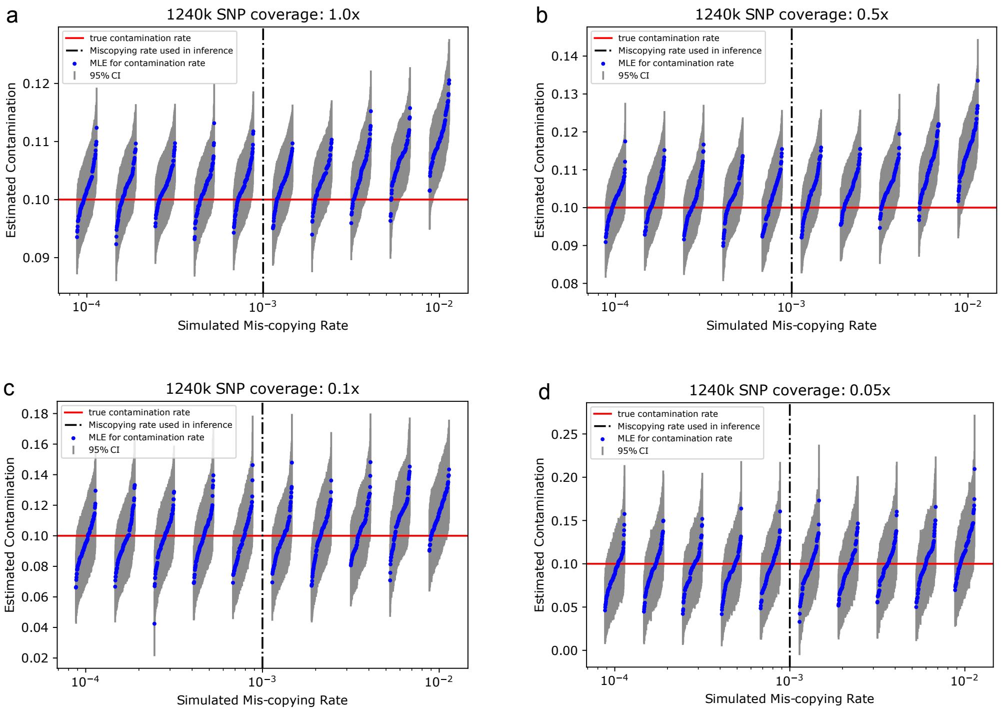


Figure S3: **Mis-specified Haplotype Copying Error Rate at Various Coverages.**

##### 1.3.3 Mis-specified Haplotype Copying Jump Rate

The haplotype copying jump rate models the rate of jumping to a different haplotype to copy from. Following Ringbauer et al. [2021], who described that $\rho = 300$ yields good performance for most modern human ancient DNA data when using Li&Stephens model for inferring ROH, we set $\rho = 300$ as the default value. We then assessed whether our model is robust with respect to a mis-specified jump rate. To do so, we simulated 10% contamination as described in Section 1.1 except that we varied the jump rate from 100 to 1000, evenly spaced on a log scale. We then estimated contamination rate when setting the jump rate to be $\rho = 300$ for all simulated samples.

The results (Fig. S4) show that the estimated contamination remains unbiased for a wide range of simulated jump rates when using the default value $\rho = 300$. Only at high jump rates close to $\rho' = 1000$, we observe some upward bias. This upward bias is minimal at relatively high coverage ($\sim$ 1x) and only at lower coverage do we start to observe moderate upward bias ($\sim$0.5x or lower, Fig. S4b,c,d). That said, previous work showed that the estimated maximum likelihood haplotype copying jump rate never exceeds 800 in 344 ancient male X chromosomes examined, with the majority of them within range 300-600 [Biddanda et al., 2021, Fig. 7]. Therefore, we believe that $\rho = 300$ is a suitable default setting that performs well on the majority of ancient DNA data.

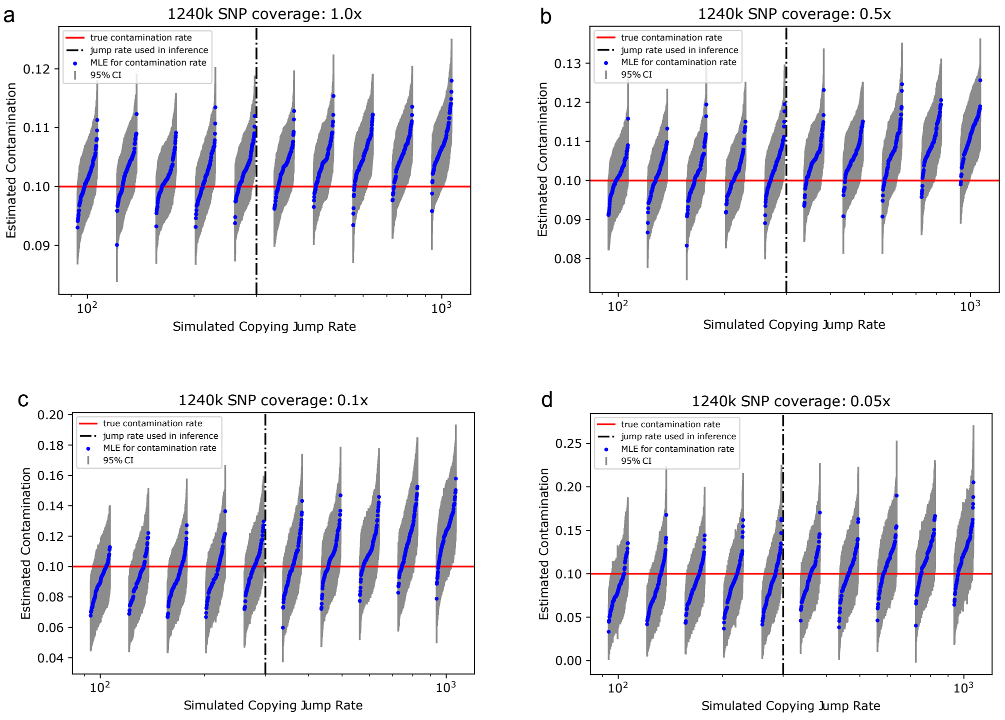


Figure S4: **Effect of Mis-specified Haplotype Copying Jump Rate at Various Coverages.**

### 2 A List of Software&Python Packages Used in This Work

- ANGSD 0.934
- samtools 1.13
- Python 3.8.10
- Numpy 1.17.4

- Scipy 1.4.1
- Numdifftools 0.9.39
- h5py 2.10.0

### 3 Supplementary Figures

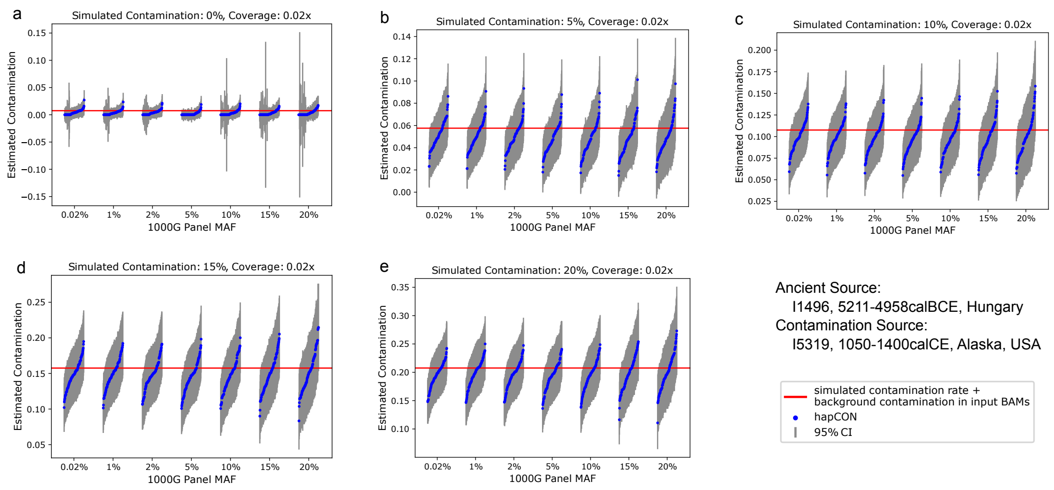


Figure S5: **Comparing hapCON on 1000G Panel with Different MAF Cutoff at Various Contamination Level.** We performed mixed BAM simulation with coverage 0.02x as described in the main article. We compared performance on the 1000Genome reference panel with varying minor allele frequency cutoffs.

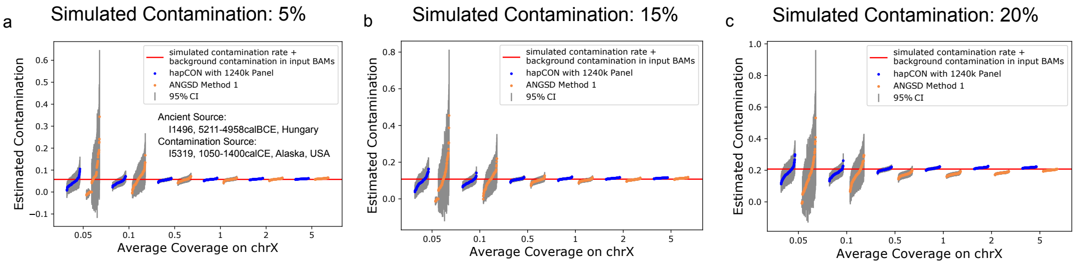


Figure S6: **Performance on 1240k Panel for Simulated Contaminated BAM Files.** We simulated contaminated BAM files by mixing two minimally contaminated BAM files. 100 replicates were taken for each simulation scenario. Baseline contamination (0.756%) from I1496 was added to the simulated contamination rate. **a** Comparing ANGSD with hapCON using 1240k panel on simulated BAM files with 5% contamination. **b** Comparing ANGSD with hapCON using 1240k panel on simulated BAM files with 15% contamination.**c**Comparing ANGSD with hapCON using 1240k panel on simulated BAM files with 20% contamination.

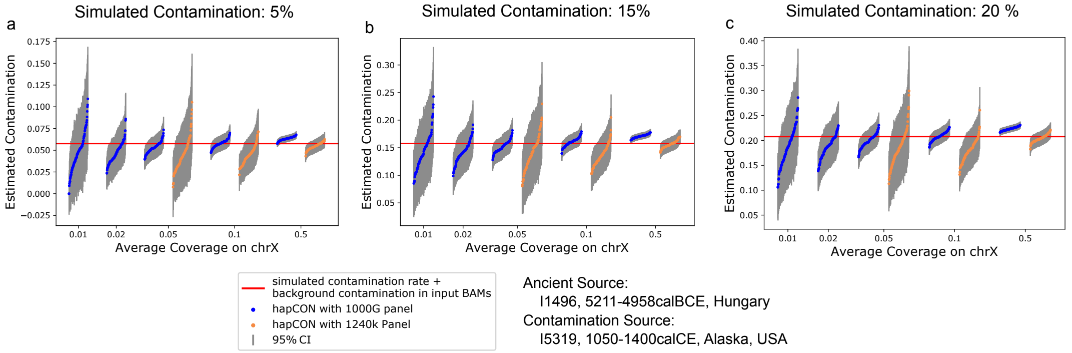


Figure S7: **Performance on 1000G Panel for Simulated Contaminated BAM Files** We compare performance on the two different reference panels: 1240k panel and 1000G panel. **a** For simulated BAM files with 5% contamination. **b** With 15% contamination. **c** With 20% contamination.

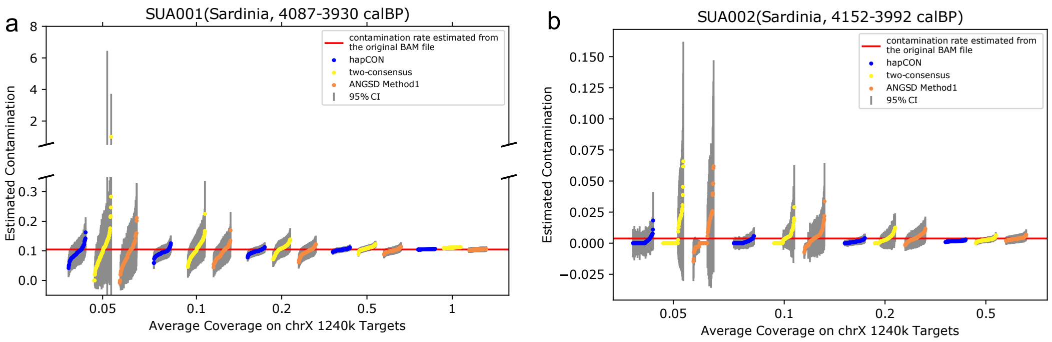


Figure S8: **Comparing hapCON, the two-consensus method and ANGSD on Downsampled Sardinia aDNA data.** Downsampling was performed as described in the main article. We compare the performance of our method, the two-consensus method and ANGSD. **a** Comparison on individual SUA001. **b** Comparison on individual SUA002.

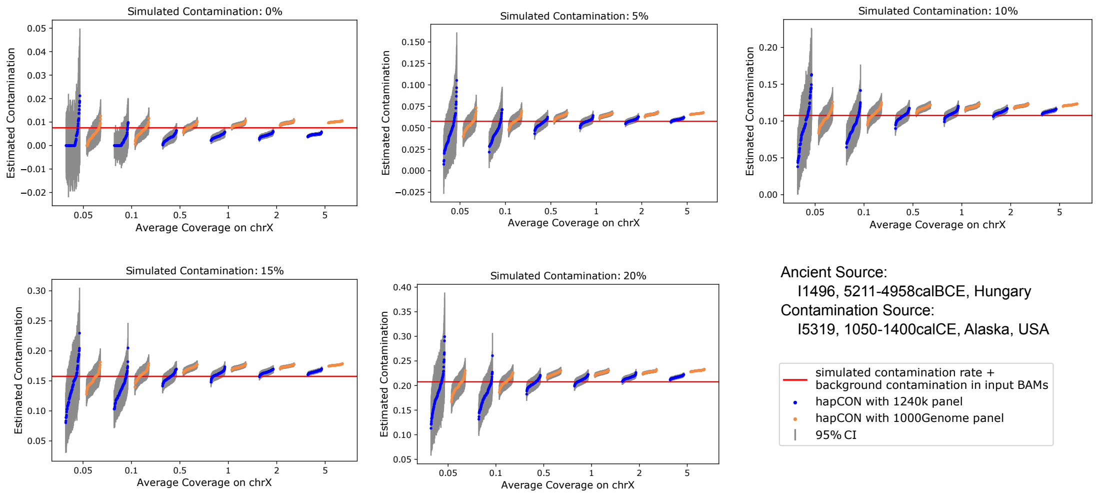


Figure S9: **Comparing hapCON at medium-to-high Coverage with Two Different Reference Panels.** We observed that hapCON tends to overestimate contamination at higher coverage. However, this bias remains mild across a wide range of coverage and contamination level. **a** 0% simulated contamination. **b** 5% simulated contamination. **c** 10% simulated contamination. **d** 15% simulated contamination. **e** 20% simulated contamination.

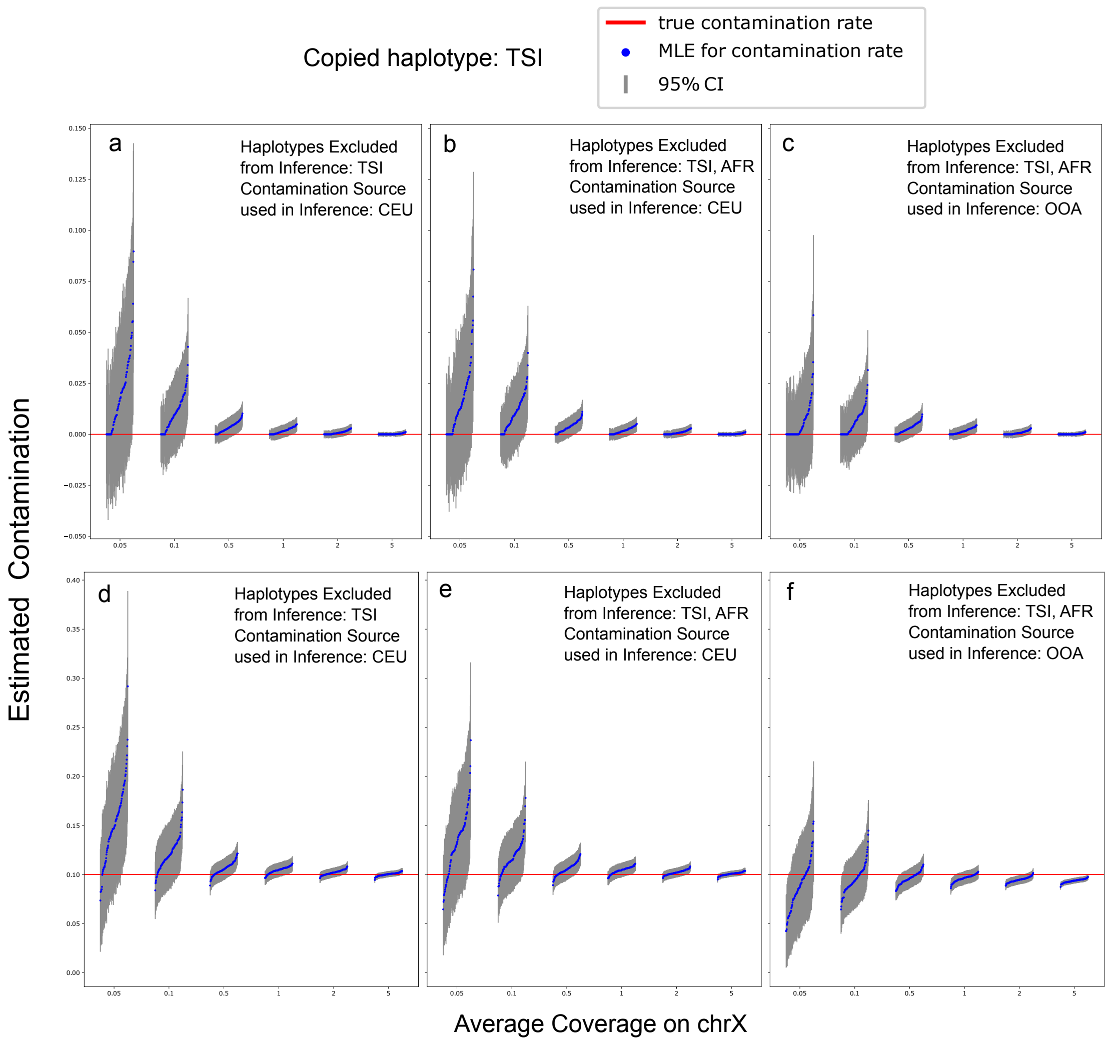


Figure S10: **Bias when the endogenous source is genetically close to the contamination source.** We performed the simulation using the default setting as described in Section 1.1 except that we used the CEU as the contamination source (rather than the global allele frequencies). Panels a-c show the results for no simulated contamination and panels d-f shows the results for simulated 10% contamination. We explored several different settings for inference by removing divergent haplotypes (AFR) from the reference panel and by using different allele frequencies as the proxy of the contamination source (CEU vs. OOA, where OOA denotes the allele frequencies of all populations in the 1000Genome except for AFR). Settings are indicated in the upper right corner of each subpanel.

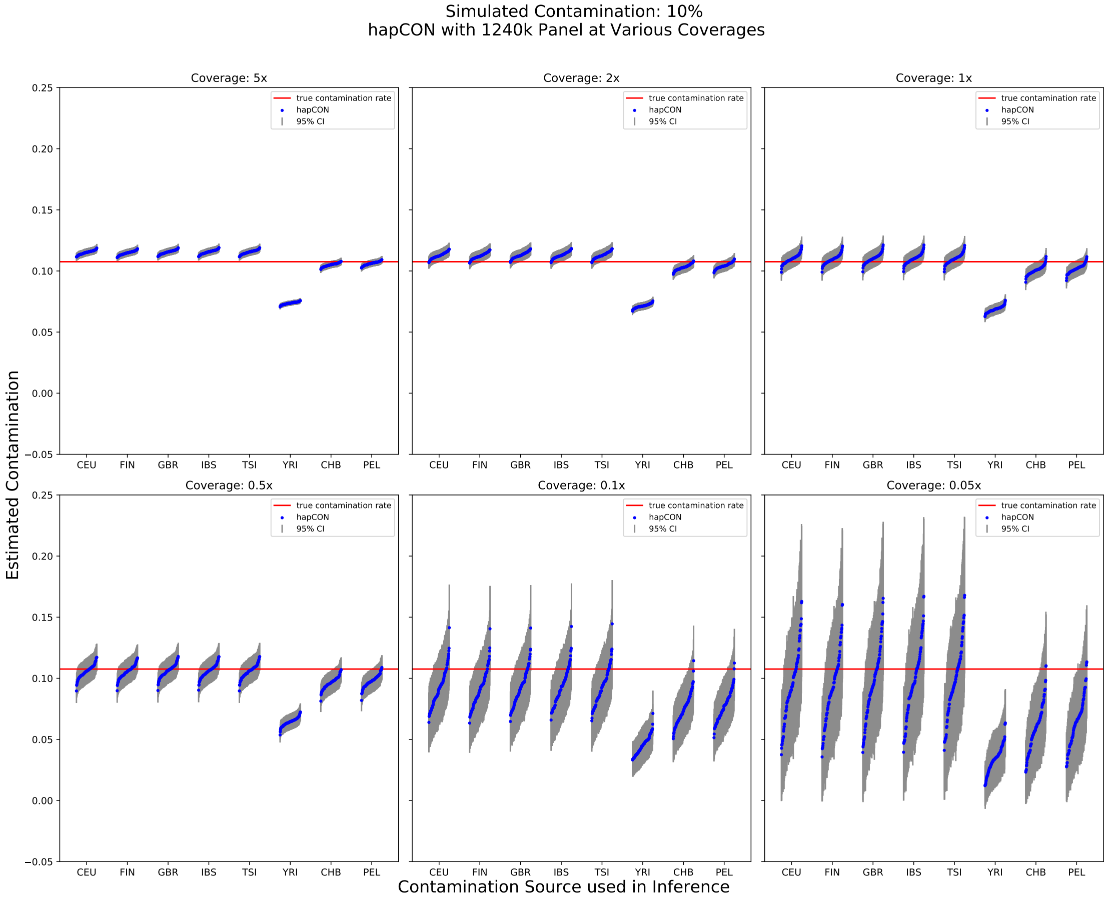


Figure S11: **Misspecified Contamination Ancestry in Mixed BAM Simulation with 1240k Panel.** We used the same mixed BAM simulation as described in section ”Simulated whole genome sequencing data” in the main article. For each coverage, we used the 1240k reference panel with CEU, FIN, GBR, IBS, TSI, YRI, CHB, PEL as the contamination ancestry to test the robustness of our method with respect to mis-specified contamination ancestry.

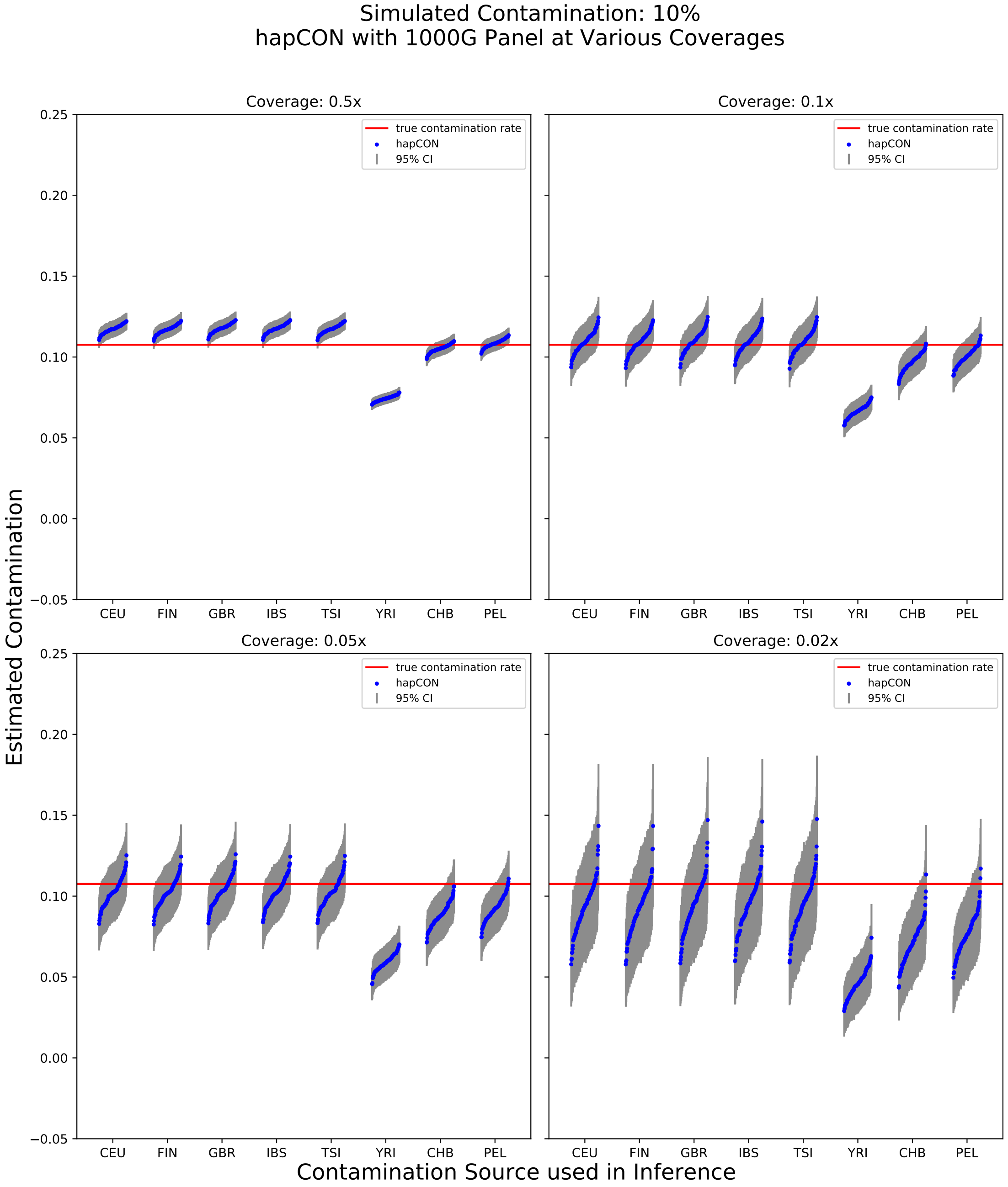


Figure S12: **Misspecified Contamination Ancestry in Mixed BAM Simulation with 1000G Panel.** Same as Fig. S11 except that we used the 1000G reference panel here.

### References

Arjun Biddanda, Matthias Steinrücken, and John Novembre. Properties of two-locus genealogies and linkage disequilibrium in temporally structured samples. *bioRxiv*, 2021.

1000 Genomes Project Consortium et al. A global reference for human genetic variation. *Nature*, 526(7571):68, 2015.

Po-Ru Loh, Petr Danecek, Pier Francesco Palamara, Christian Fuchsberger, Yakir A Reshef, Hilary K Finucane, Sebastian Schoenherr, Lukas Forer, Shane McCarthy, Goncalo R Abecasis, et al. Reference-based phasing using the haplotype reference consortium panel. *Nature Genetics*, 48(11):1443–1448, 2016.

J Víctor Moreno-Mayar, Thorfinn Sand Korneliussen, Jyoti Dalal, Gabriel Renaud, Anders Albrechtsen, Rasmus Nielsen, and Anna-Sapfo Malaspinas. A likelihood method for estimating present-day human contamination in ancient male samples using low-depth X-chromosome data. *Bioinformatics*, 36(3):828–841, 2020.

Iñigo Olalde, Swapan Mallick, Nick Patterson, Nadin Rohland, Vanessa Villalba-Mouco, Marina Silva, Katharina Dulias, Ceiridwen J Edwards, Francesca Gandini, Maria Pala, et al. The genomic history of the Iberian Peninsula over the past 8000 years. *Science*, 363(6432):1230–1234, 2019.

Morten Rasmussen, Xiaosen Guo, Yong Wang, Kirk E Lohmueller, Simon Rasmussen, Anders Albrechtsen, Line Skotte, Stinus Lindgreen, Mait Metspalu, Thibaut Jombart, et al. An Aboriginal Australian genome reveals separate human dispersals into asia. *Science*, 334(6052):94–98, 2011.

Harald Ringbauer, John Novembre, and Matthias Steinrücken. Parental relatedness through time revealed by runs of homozygosity in ancient DNA. *Nature Communications*, 12(1):1–11, 2021.

Simone Rubinacci, Diogo M Ribeiro, Robin J Hofmeister, and Olivier Delaneau. Efficient phasing and imputation of low-coverage sequencing data using large reference panels. *Nature Genetics*, 53(1):120–126, 2021.
